# Responses of Neurons in the Medullary Lateral Tegmental Field and Nucleus Tractus Solitarius to Vestibular Stimuli in Conscious Felines

**DOI:** 10.1101/2020.10.23.352542

**Authors:** John P. Bielanin, Nerone O. Douglas, Jonathan A. Shulgach, Andrew A. McCall, Derek M. Miller, Pooja R. Amin, Charles P. Murphey, Susan M. Barman, Bill J. Yates

## Abstract

Considerable evidence shows that the vestibular system contributes to adjusting sympathetic nervous system activity to maintain adequate blood pressure during movement and changes in posture. However, only a few prior experiments entailed recordings in conscious animals from brainstem neurons presumed to convey baroreceptor and vestibular inputs to neurons in the rostral ventrolateral medulla (RVLM) that provide inputs to sympathetic preganglionic neurons in the spinal cord. In this study, recordings were made in conscious felines from neurons in the medullary lateral tegmental field (LTF) and nucleus tractus solitarius (NTS) identified as regulating sympathetic nervous system activity by exhibiting changes in firing rate related to the cardiac cycle, or cardiac-related activity (CRA). Approximately 38% of LTF and NTS neurons responded to static 40° head up tilts with a change in firing rate of ^~^50%. However, few of these neurons responded to 10° sinusoidal rotations in the pitch plane, in contrast to prior findings in decerebrate animals that the firing rates of both NTS and LTF neurons are modulated by small-amplitude body rotations. Thus, as previously demonstrated for RVLM neurons, in conscious animals NTS and LTF neurons only respond to large rotations that lead to changes in sympathetic nervous system activity. The similar responses to head-up rotations of LTF and NTS neurons with those documented for RVLM neurons suggest that LTF and NTS neurons are components of the vestibulo-sympathetic reflex pathway. However, a difference between NTS/LTF neurons and RVLM was variability in CRA over time. This variability was significantly greater for RVLM neurons, raising the hypothesis that the responsiveness of these neurons to baroreceptor input is adjusted based on the animal’s vigilance and alertness.

## 1 Introduction

The neural pathways through which baroreceptor and other inputs affect sympathetic nervous system activity have been determined in a variety of mammalian species [1; 2; 3]. A caveat is that most neurophysiologic experiments examining changes in activity of neurons in key brainstem areas that participate in regulating cardiovascular function were conducted in anesthetized or decerebrate animals [4]. An exception is the rostral ventrolateral medulla (RVLM), an area of the reticular formation that plays a key role in controlling the firing rate of sympathetic preganglionic neurons that regulate heart rate and constriction of vascular smooth muscle [5; 6]. Three studies reported the activity of RVLM neurons in conscious felines and how it differs from that in decerebrate animals [7; 8; 9]. These studies showed that RVLM neurons whose activity is correlated with the cardiac cycle, and presumably participate in controlling sympathetic nervous system activity, have spontaneous firing rates that vary over time. They also compared peak and trough firing rates of RVLM neurons between cardiac cycles and determined that they can also fluctuate during a recording session. It was postulated that these changes in spontaneous firing rate and cardiac-related activity (CRA) of RVLM neurons were dependent on supratentorial inputs and related to the animal’s alertness, vigilance, and attentiveness to its environment [4; 7; 9].

Studies have shown that much larger changes in body position are required to modulate the activity of RVLM neurons in conscious animals than in decerebrate animals [8]. Inputs from the vestibular system participate in regulating sympathetic nervous system activity during changes in body position [10]. In decerebrate felines, vestibular inputs elicited by small (<10°) tilts were sufficient to alter the activity of sympathetic efferent fibers in peripheral nerves as well as the firing rate of RVLM neurons [8; 11]. However, in conscious animals the firing rates of RVLM neurons were unaffected by 10° sinusoidal body rotations [8; 9], although the activity of a third of the neurons was modulated by static 40° head-up tilts [9]. These changes in unit activity were not correlated with changes in heart rate, suggesting that they were not due to engagement of the baroreceptor reflex. Gating of vestibular inputs to RVLM neurons such that these neurons only respond to large-amplitude static tilts has physiologic relevance, as head-up tilts <40° in amplitude do not appreciably affect the distribution of blood in the body requiring a compensatory change in sympathetic nervous system activity [12]. This is unlike other vestibular-elicited responses, such as vestibulo-ocular reflexes, which are engaged during changes in head position less than 1° in magnitude [13].

Baroreceptor inputs are relayed to RVLM neurons through a pathway that includes neurons in nucleus tractus solitarius (NTS), which receive direct inputs from baroreceptor afferents [1; 2; 3]. NTS neurons relay baroreceptor signals to the RVLM through connections in the medullary reticular formation [1; 2; 3]. In felines, neurons in the dorsolateral medullary reticular formation, a region termed the lateral tegmental field (LTF), also play a key role in relaying baroreceptor signals to the RVLM [14; 15; 16; 17]. There is also evidence that both NTS and LTF neurons transmit vestibular signals to RVLM neurons. NTS [18; 19; 20] and LTF [21] receive direct projections from the vestibular nuclei, and experiments in decerebrate animals showed that NTS [22] and LTF [23] neurons respond to small-amplitude rotations of the animal’s body. To our knowledge, the activity of NTS neurons with baroreceptor inputs has not been recorded in conscious animals. Although one prior study considered the responses of LTF neurons to vestibular stimuli in conscious animals, the focus of this work was neurons that lacked CRA that presumably were part of the “vomiting center,” which also is located in the dorsolateral medullary reticular formation [24].

The present study examined CRA and responses to 10° sinusoidal rotations and 40° static tilts of NTS and LTF neurons in conscious felines. The major goal was to ascertain whether NTS and LTF neurons are likely components of the vestibulo-sympathetic pathway in conscious animals, with responses to vestibular stimuli similar to those of RVLM neurons presumed to transmit these signals to sympathetic preganglionic neurons in the spinal cord [9]. In addition, comparing the responses to whole-body tilts of NTS and LTF neurons with those of RLVM neurons provides insights into the role of neurons at each stage of the vestibulo-sympathetic reflex pathway in transforming signals from the inner ear. As noted above, changes in sympathetic nervous system activity are elicited only during large-amplitude changes in body position that can result in decreased venous return to the heart and diminished cardiac output [4; 12]. A gating of signals thus occurs in the vestibulo-sympathetic circuitry to suppress responses to vestibular inputs elicited during small-amplitude movements. It is informative to compare how NTS and LTF neurons respond to small- and large-amplitude rotations in conscious animals, to determine if they are components of this gating mechanism.

A secondary goal of these experiments was to establish whether anticipation of a passive movement elicits changes in NTS and LTF neuronal activity. It is well-established that feedforward cardiovascular responses are elicited prior to exercise and some other active movements [25; 26; 27]. A study that monitored changes in heart rate and cerebral blood flow in conscious felines before and during 60° head-up tilts preceded by a light cue demonstrated that no preparatory cardiovascular responses occur before such imposed changes in body position [28]. The conclusion of this study was that preparatory cardiovascular responses occur before active movements, but not passive (imposed) movements requiring increases in sympathetic nervous system activity. This conclusion was supported by a study showing that no changes in RVLM neuronal activity occurred in conscious animals following a light cue and before the onset of head-up tilts [9]. However, it is possible that changes in neuronal activity are present upstream in the vestibulo-sympathetic reflex pathway, but suppressed prior to being relayed to the RVLM. Thus, by determining whether the activity of LTF and NTS neurons changes preceding cued head-up tilts, the present study provided an additional test of the hypothesis that feedforward cardiovascular responses occur only before active movements, but are absent prior to expected passive changes in body position [28].

## 2 Methods and Materials

All experimental procedures on animals followed the National Research Council’s *Guide for the Care and Use of Laboratory Animals* [29] and were approved by the University of Pittsburgh’s Institutional Animal Care and Use Committee. Experiments were performed on five female antibody profile defined and specific pathogen free (APD/SPF) domestic shorthair cats obtained from Marshall BioResources (North Rose, New York, USA). Animals were 4-6 months of age when acquired from the vendor. Juvenile female animals were obtained for these studies, following our prior experience that they are more amenable than males to be acclimated for the 2-hour restraint period required for data collection. Moreover, the much higher growth rate in the first year for male cats makes it difficult to maintain chronically implanted recording devices. Animals were provided commercial cat food and water *ad libitum* and were housed under 12-h light/dark cycles. Table 1 provides information about the animals and the parameters of the study.

**Table 1.** Information about the animals used in the study, the length of data collection, and the number of units included in data analysis.

| Animal | Weight at Surgery (kg) | Total period of data collection (days) | LTF Units ≥ 5 Trials | | NTS Units ≥ 5 Trials | |
| --- | --- | --- | --- | --- | --- | --- |
|  |  |  | CRA | No CRA | CRA | No CRA |
| 1 | 3.4 | 266 | 5 | 1 | 0 | 0 |
| 2 | 3.4 | 211 | 8 | 8 | 3 | 3 |
| 3 | 4.2 | 399 | 9 | 2 | 17 | 1 |
| 4 | 4.7 | 213 | 6 | 2 | 1 | 0 |
| 5 | 4.2 | 112 | 4 | 0 | 0 | 0 |
| Total Number of Units Included in Analysis |  |  | 32 | 13 | 21 | 4 |

This study employed procedures similar to those used in our prior experiments [7; 8; 9; 28; 30], which entailed acclimating animals for restraint on a computer-controlled tilt table, implanting instrumentation for recording the electrocardiogram (ECG) and single unit neuronal activity from the brainstem, and subsequently collecting data over a period of 112-399 days. Following data collection, animals were euthanized, and unit locations were histologically reconstructed.

### Surgical Procedures

To avoid changes in hormone levels that could affect physiological responses, animals were spayed by a veterinarian prior to the beginning of the study. After they were spayed, the animals were conditioned for body restraint in an animal holder mounted on a computer-controlled tilt table. The period of gradual acclimatization for restraint lasted about 2 months per animal but varied based on the animal’s behavior. At the beginning of acclimatization, animals were initially restrained for only a few minutes per session, but as training continued, the period of restraint was slowly increased to two hours.

After an animal was acclimated to 2 hours of body restraint, a recovery surgery was performed in a dedicated operating suite using aseptic techniques. The recovery surgery entailed a craniectomy, attachment of a head fixation plate and recording chamber to the skull, and implantation of wire bundles beneath the thoracic skin to record the ECG. These procedures were described in detail in prior manuscripts [7; 8; 9; 28; 30]. During surgeries, animals were initially anesthetized by an intramuscular injection of ketamine (20 mg/kg) and acepromazine (0.2mg/kg), and anesthesia was maintained using 1-2% isoflurane vaporized in O2 provided through an endotracheal tube. Vital signs (heart rate, respiration rate, and blood oxygen saturation) were measured and recorded in the animal’s anesthesia record every 15 minutes. Ringer lactate solution was provided intravenously and a heating lamp and pad were used to maintain body temperature near 38°C. Following the surgery, an antibiotic (amoxicillin, 50mg/kg, orally twice per day) was provided for 10 days, and a transdermal fentanyl patch (25 μg/hr) delivered analgesia for 72 hours.

### Recording Procedures

After 2 weeks of recovery from surgery, animals were acclimated for a month to head restraint by inserting a screw into the head-mounted fixation plate, and 40° whole-body tilts in the pitch plane provided by a servo-controlled hydraulic tilt table (Neurokinetics, Pittsburgh, PA USA), which was operated using a Micro1401 mk 2 data collection system and Spike-2 version 7 software (Cambridge Electronic Design, Cambridge, UK). During experiments, the recording room was darkened, and black curtains were placed around the tilt table to mask any visual cues about the animal’s position in space. An array of 10 LED lights generating a light intensity of 300 lumens was positioned 28 cm in front of the animal’s face to provide a visual prompt prior to tilts. The light array was controlled by the Cambridge Electronic Design hardware and software, and illuminated for 2 seconds, from 8-10 seconds prior to the onset of tilts (see Fig. 1). The light cue was presumably highly salient since the area surrounding the animal was darkened. The light cue was provided during every trial during the acclimation period, so animals had experienced several hundred tilts paired with a light cue prior to the onset of recordings. Animals were continuously monitored during recording sessions for indicators of distress, and to assure that they remained awake with eyes open.

**Fig. 1.**
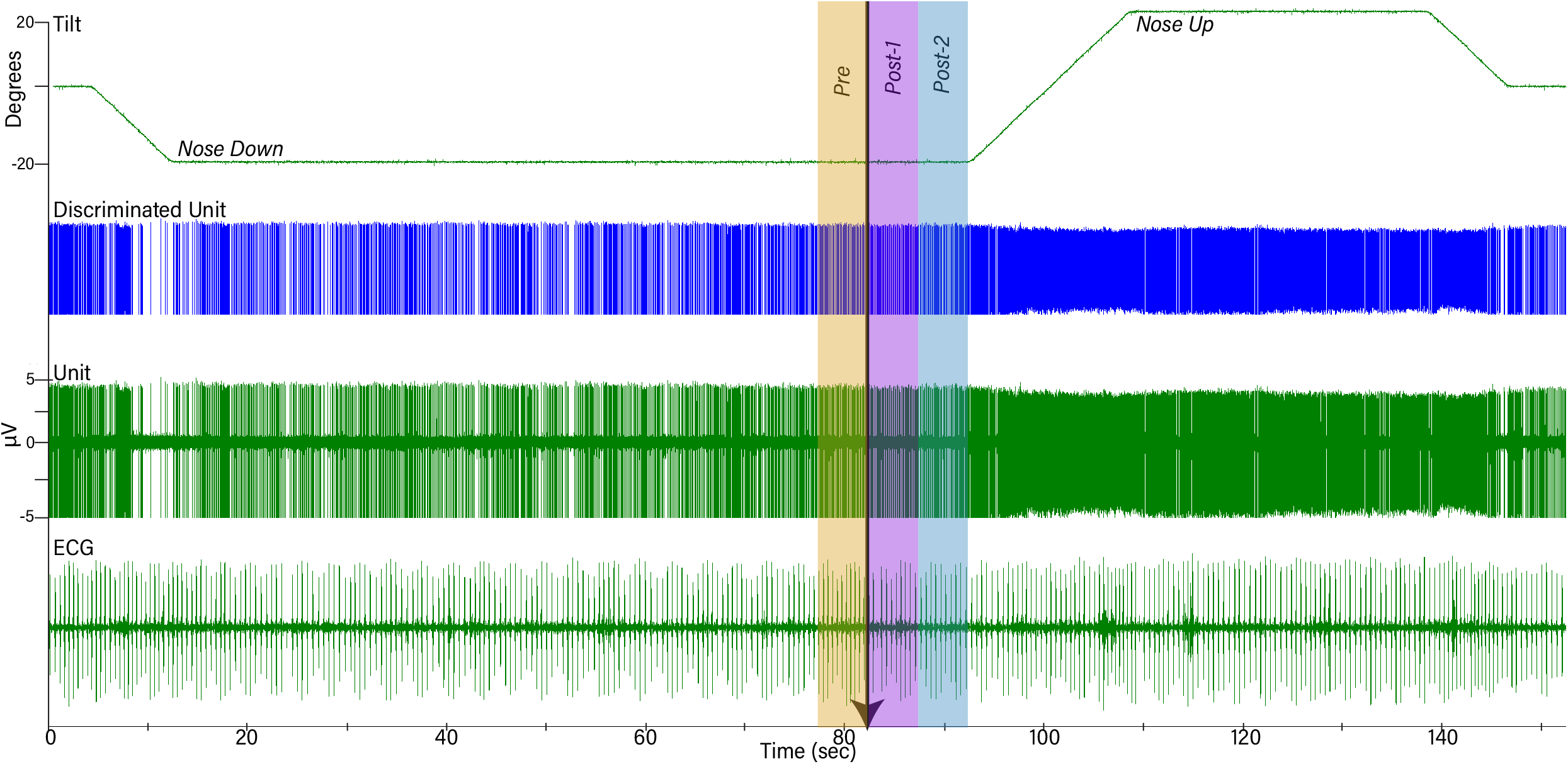
Example of data collected during one trial. The top trace shows table position, the second shows discriminated activity for one unit, the third shows raw unit activity recorded by a microelectrode, and the bottom trace is a recording of the electrocardiogram (ECG). Initially, animals were positioned 20° nose-down for a variable amount of time. A light cue was presented (onset indicated by arrow) that lasted for two seconds. Ten seconds after the light cue was initiated, animals were tilted 40° head-up (20° nose-down to 20° nose-up) and remained in that position for 30 seconds prior to returning to the prone position. To determine if the light cue had any effect on a unit’s activity, firing rate during the period five seconds before light onset (Pre-) was compared to that during the period 5-10 seconds prior to tilt onset (Post-2).

Brainstem neuronal recordings were initiated once the animals were acclimated for restraint on the tilt table. A hydraulic microdrive (model 650, David Kopf Instruments, Tujunga, CA USA) and x-y positioner, both attached to the recording chamber, allowed for epoxy-insulated tungsten microelectrodes (5MΩ, Frederick Haer, Bowdoin, ME USA) to be inserted at defined coordinates into the brainstem. Recordings were performed over a 122-399 day period (see Table 1) and targeted to LTF and NTS neurons located 0-5 mm rostral to the obex. Units were localized using stereotaxic coordinates and physiological landmarks including the dorsal and ventral respiratory groups (which were evident as clusters of neurons with respiratory-related activity).

Activity recorded from brainstem neurons was amplified by a factor of 10,000 and filtered with a band pass of 300-10,000 Hz using an AM Systems (Sequim, WA USA) model 1800 microelectrode AC amplifier. The output of the amplifier was sampled at 25,000 Hz using the Micro1401 mk 2 data collection system and Spike-2 version 7 software. The ECG signal was amplified by a factor of 1,000 and filtered with a bandpass of 10-10,000 Hz using an AM Systems model 1700 differential AC amplifier and sampled at 2,500 Hz. Voltages from potentiometers mounted on the tilt table that signaled table position were sampled at 100 Hz. An example of data recordings is shown in Fig. 1.

When a presumed NTS or LTF neuron was isolated, the animal was tilted 20° nose-down. The animal remained in this position for 1-4 minutes to allow baseline recordings to be collected to determine if the cell exhibited cardiac-related activity (CRA). Subsequently, the light cue was provided, and the animal was tilted 20° nose-up (such that the table movement was 40°). Since the period when the animal was tilted nose-down varied between trials, animals were not able to predict the occurrence of the light cue or the following nose-up tilt. After the animals were tilted nose-up, they remained in this position for 30 seconds prior to returning to their initial prone position (see Fig. 1). Multiple trials (up to 13) were conducted if unit activity remained stable. For most units responses to 10° (peak-to-peak) 0.5 Hz sinusoidal rotations in the pitch plane were also recorded.

### Data Analysis Procedures

Typically, only one or two units were present in the recording field during trials. The spike detection and sorting algorithm of the Spike-2 software was used to isolate the activity of each unit in the recording field, and to verify that spike shape remained consistent between trials. If there was any doubt that data were not continually recorded from the same unit, the data were discarded. Event markers generated by the Spike-2 software indicated the timing of action potentials generated by each sampled neuron, as well as the peak ECG R-waves. These event markers were used in our subsequent data analyses.

#### Detection of CRA

Experimental procedures similar to those used in our prior studies were used to determine if a unit exhibited CRA [7; 8; 9]. Trigger pulses coinciding with the R-wave of the ECG were used for the creation of post R-wave interval histograms (10-ms bin width) of unit activity (Datapac software, Run Technologies; Mission Viejo CA USA). Interval histograms between successive R-waves were constructed simultaneously. Trigger pulses were also used to generate an average of the peak ECG R-wave. The averages and histograms were generated using data segments of varying length (28 – 211 s). When multiple trials were conducted for each neuron, a separate analysis was performed for each trial. A ratio of peak-to-background counts was calculated for each histogram. To classify a neuron as having CRA, the ratio of peak-to-background counts in the histogram had to be greater than a value of 2.0 for the majority of analyses for each unit. Figure 2 shows examples of two analyses for an NTS unit.

**Fig. 2.**
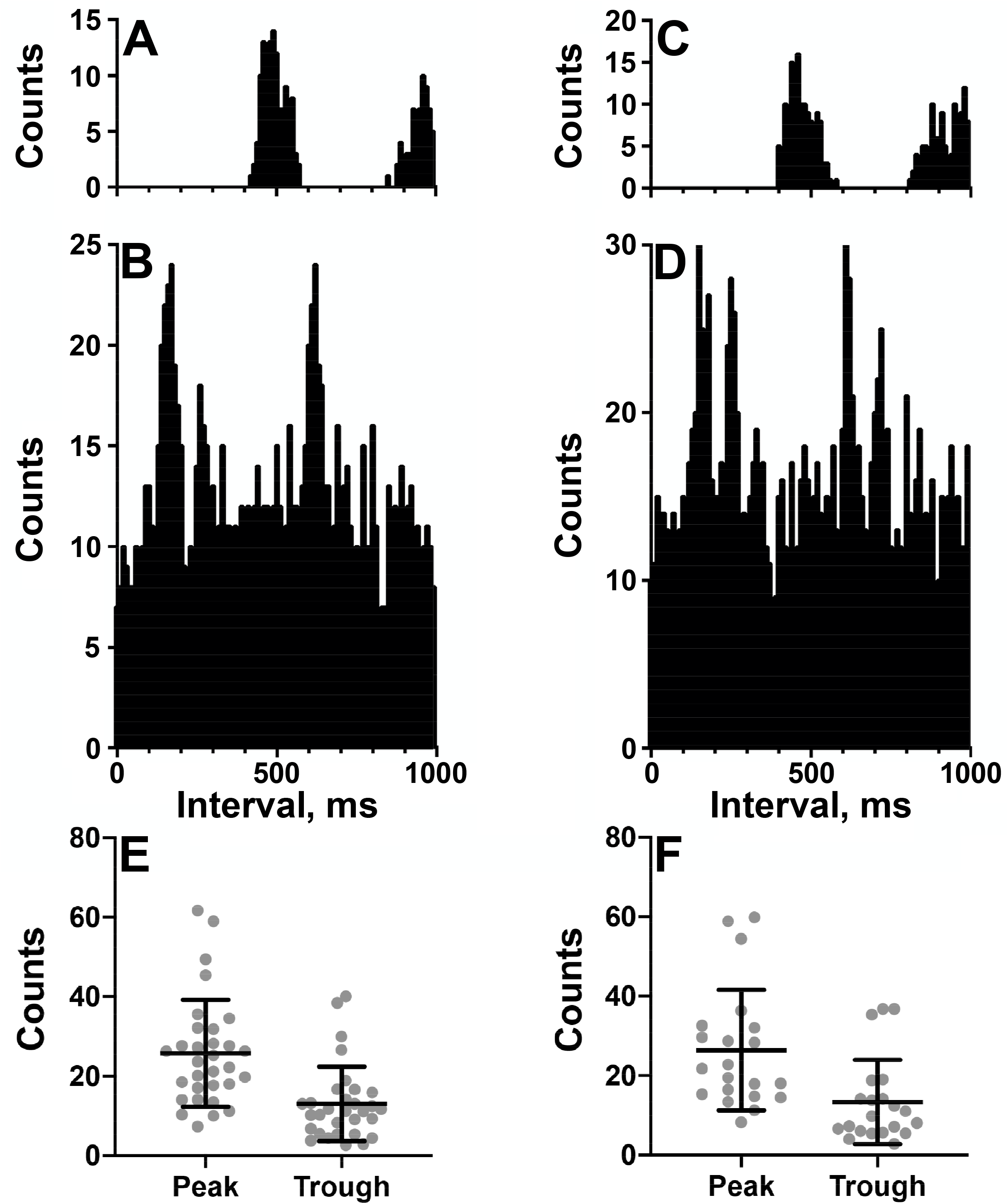
*A-D:* Cardiac-related activity (CRA) evident in two trials for an NTS neuron (*A* and *B* are for one trial and *C* and *D* are for the second trial). Histograms were triggered by event markers demarking ECG R-waves; bin width is 10 ms. *A* and *C* show intervals between R-waves, and *B* and *D* show intervals between the unit’s action potentials. Unit activity peaked in periods between R-waves. *E-F*: plot of counts during peak and trough periods for the LTF (*E*) and NTS (*F*) neurons classified as exhibiting CRA.

#### Response of Units to Light Cue and 40° Tilts

The following analyses were conducted using MATLAB software (Mathworks, Natick, MA USA). Unit firing rate and heart rate were determined for a 30-sec period prior to the light cue, while the animal was tilted nose-down, as well as when the animal was positioned nose-up. Furthermore, unit firing rate and heart rate were also determined for the time period 5 seconds prior to the light cue and during the two consecutive 5-second time periods prior to the onset of the 40° nose-up tilt (see Fig. 1). Since the duration of the light cue was 2 seconds, the light remained illuminated during the first 2 seconds of Post-1 time interval indicated in Fig. 1.

Subsequent statistical analysis of the data compared heart rate and unit firing rates when animals were positioned nose-up and nose-down, as well as during the pre- and post-light cue periods prior to the onset of the tilt. As established in a prior study [9], to provide adequate statistical power these analyses were only conducted for units in which five or more 40° tilts were delivered. Statistical analyses and plotting of experimental results were performed using Prism 8 software (GraphPad software, San Diego, CA USA). Confidence intervals are indicated as mean ± one standard deviation.

#### Response to Sinusoidal Rotations

We additionally tested the responses of most neurons to 10° (peak-to-peak amplitude) sinusoidal tilts at 0.5 Hz to determine if they responded to small dynamic vestibular stimuli as well as large amplitude tilts. During sinusoidal rotations, neural activity was binned (500 bins/cycle) for ^~^ 35 stimulus repetitions and fitted with a sine wave using a least-squares minimization technique described in other reports [8; 22; 24; 30; 31]. Two criteria were used to determine if a unit’s activity was significantly modulated by 10° sinusoidal rotations: a signal-to-noise ratio > 0.5 and only one evident first harmonic. To ensure that a unit’s response to the rotations was not a reflection of its rhythmic spontaneous firing rate at the stimulus frequency, the stimulus was periodically stopped and restarted to determine if peak firing rate remained aligned with the same phase of head movement. Units were discarded from analysis if they were determined to have rhythmic activity at the frequency of the stimulus, although such occurrences were rare.

#### Reconstruction of Unit Locations

Once data collection was completed in an animal, electrolytic lesions were made at designated coordinates by passing a 100-μA negative current for 60 seconds through a 0.5-MΩ tungsten electrode. Approximately 1 week following the lesions, animals were anesthetized by an intramuscular injection of ketamine (20 mg/kg) and acepromazine (0.2 mg/kg) followed by an intraperitoneal injection of pentobarbital sodium (40 mg/kg). Animals were then transcardially perfused with saline followed by 10% formalin. The brain was removed and placed in a solution of 10% formalin / 30% sucrose for at least two weeks to allow for additional fixation. After the brain was post-fixed, it was sectioned transversely at 50-μm thickness using a freezing microtome. The sections were mounted in serial order on microscope slides and stained using thionine. Images of sections were captured using a spot camera and digital software (Spot Imaging, Sterling heights, MI USA) through a Nikon Eclipse E600 microscope. Photomontages of sections were created with Adobe Illustrator software (Adobe Inc., Mountain View, CA USA) and used to plot specific unit locations. The locations of individual units were reconstructed with respect to the position of the lesions, the relative coordinates of each unit and unit depths.

## 3 Results

Activity was recorded from a total of 134 units with CRA whose locations were histologically confirmed to be in the LTF (97 neurons) and NTS (57 neurons). The locations of the units are shown in Fig. 3. Additionally, unit activity was recorded from 51 units without CRA: 39 neurons in the LTF and 12 neurons in the NTS. Thus, 71% of the sampled LTF neurons and 83% of the sampled NTS neurons had CRA. Recordings from LTF neurons occurred in all experiments, whereas recordings of NTS activity were included in three experiments.

**Fig. 3.**
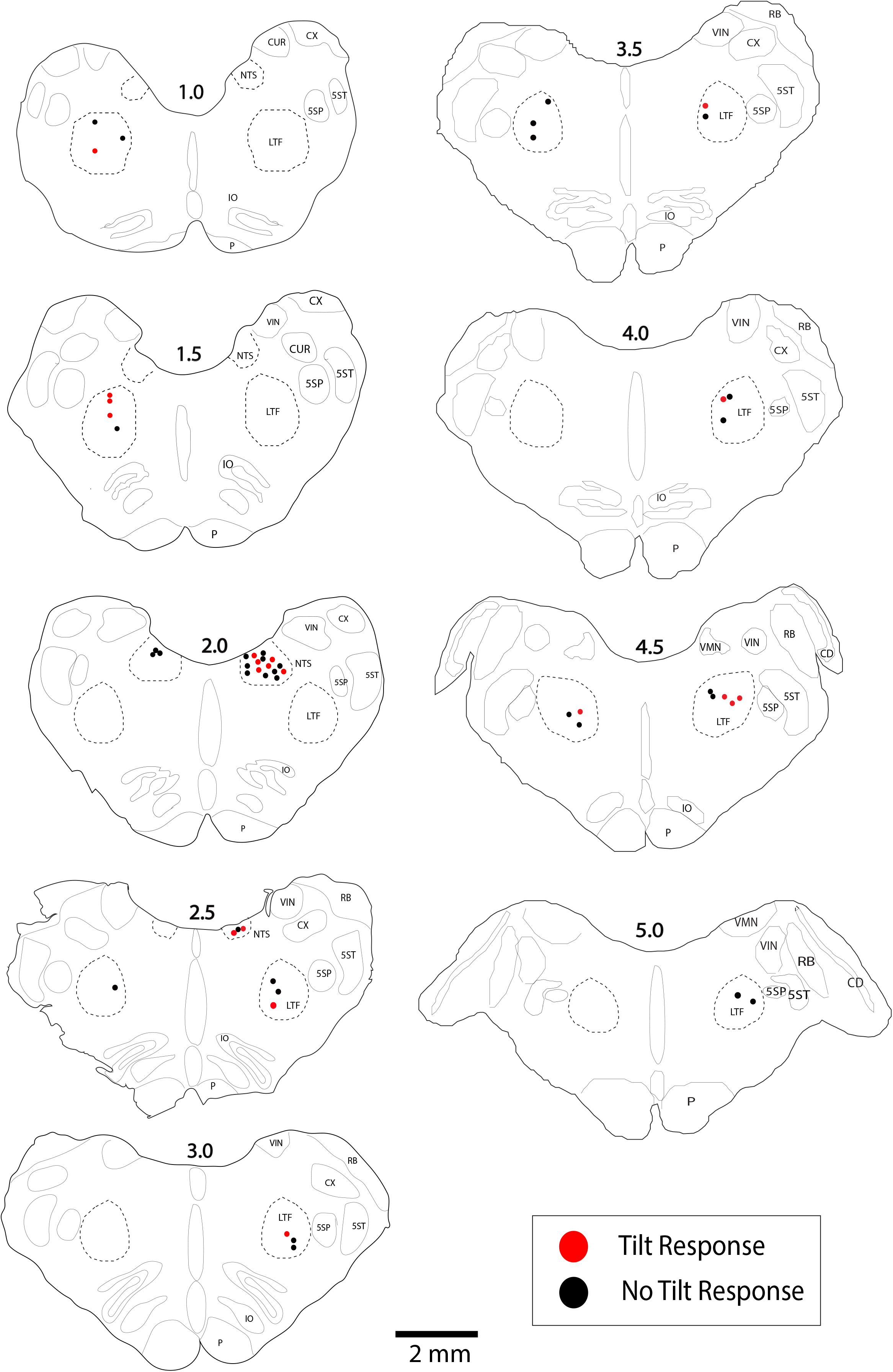
Locations of LTF and NTS neurons with CRA whose responses to 40° tilts were recorded during five or more trials. Red symbols designate units with significant differences in unit activity in the nose-down and nose-up position. Black symbols designate units without significant responses to tilts. The numbers above each section indicate the distance rostral to the obex in mm. Abbreviations: *LTF*, lateral tegmental field; *NTS*, nucleus tractus solitarius; *5SN*, spinal trigeminal nucleus; 5SŢ, spinal trigeminal tract; *CUR*, cuneate nucleus, rostral; *CX*, external cuneate nucleus; *IO*, inferior olivary nucleus; *P*, pyramid; *VIN*, inferior vestibular nucleus; *RB*, restiform body; *VMN*, medial vestibular nucleus; *CD*, dorsal cochlear nucleus.

Figure 2 A-D shows examples of CRA in two trials for an NTS unit; Fig. 2E compares the peak and trough activity for LTF neurons with CRA, and Fig. 2F provides the same comparison for NTS neurons. Each point in Figs. 2E-F is an average of all of the trials conducted for a cell. For every unit examined with five or more trials, a two-tailed paired t-test showed that on average peak counts were significantly greater (P<0.05) than trough counts. For units located in the LTF, P values ranged from <0.0001 to 0.02, with most units having P<0.001, and only 5 having P values between 0.002 and 0.02. On average, peak counts were 2.2±0.5 (SD) times greater than trough counts. Similarly, for NTS units with five or more stimulus trials, P values ranged from <0.0001 to 0.01, with most units having P<0.0004, and only 7 having P values between 0.001 and 0.01. On average, peak counts were 2.2±0.4 (SD) times greater than trough counts. Even though CRA was present across trials for units that demonstrated this activity, the ratio for peak to trough counts often varied between trials, as illustrated in Fig. 4 for eight units each in the LTF and NTS. For LTF units, the difference in peak-to-trough ratios for a particular unit (the largest ratio minus the smallest ratio across trials) ranged from 0.7 to 2.7; the median difference was 1.0. For NTS units, the difference in peak-to-trough ratios ranged from 0.3 to 2.9; the median difference was also 1.0. The variability in peak-to-trough ratios between runs was not significantly different for LTF and NTS neurons (P=0.84, unpaired two-tailed t-test).

**Fig. 4.**
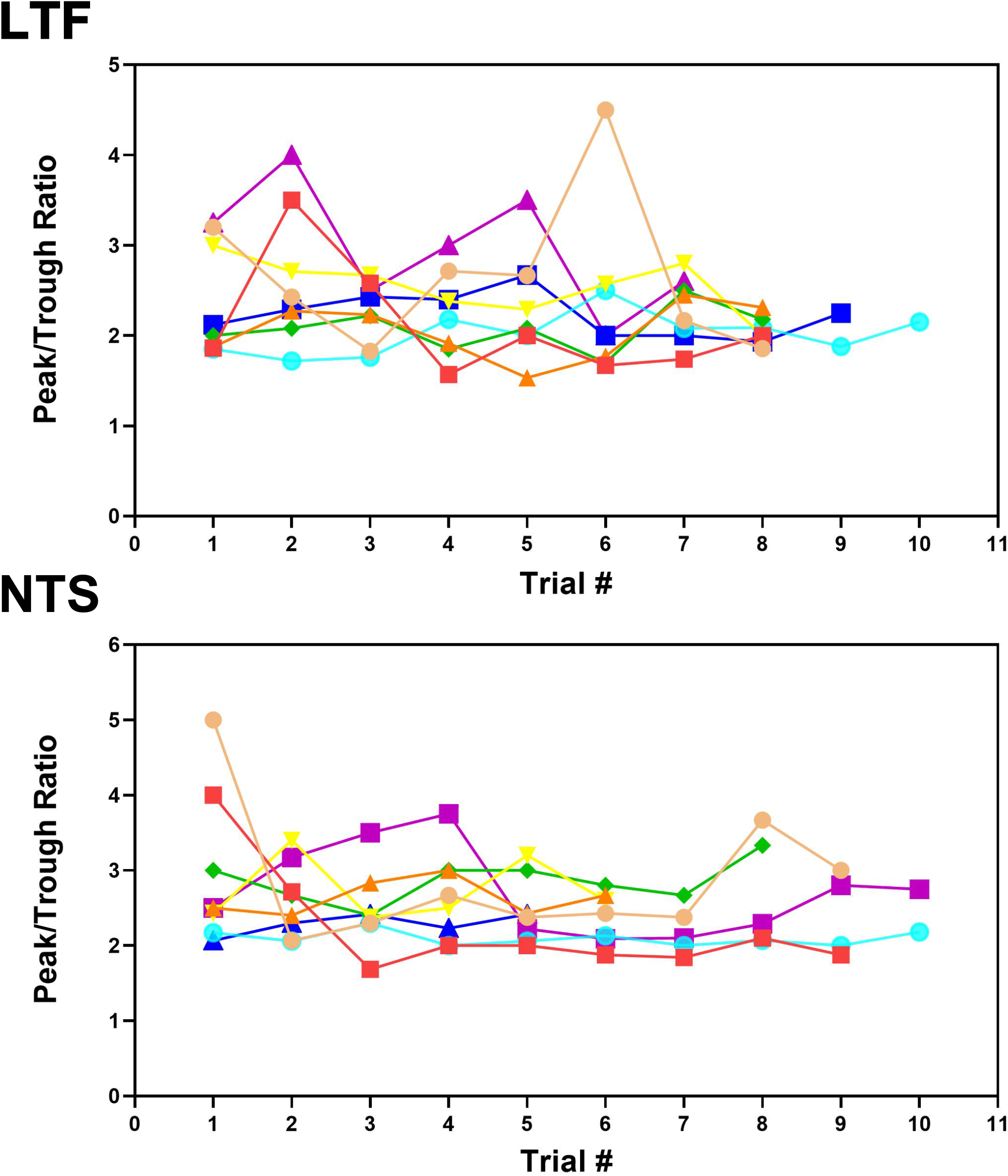
Peak-to-trough ratio of activity related to the cardiac cycle for 8 LTF and 8 NTS units where multiple trials were conducted. Data for each unit is depicted by different shapes and colors. For most units, the peak-to-trough ratio varied throughout the course of recording although all units exhibited CRA during every trial.

A similar analysis for RVLM data [9] showed that the variability in peak-to-trough ratios ranged from 2.8 to 4.3; the median difference was 3.4 (mean of 3.5 ± 0.6). A one-way ANOVA combined with Dunnett’s post-hoc test showed that the variability in the peak-to-trough ratios was significantly larger for RVLM neurons than either NTS or LTF neurons (P<0.0001). This variability in peak-to-trough ratios did not appear to be related to unit spontaneous firing rate, which was not significantly different (p=0.15, one-way ANOVA) for LTF (14±6 spikes/s), NTS (11±6 spikes/s), and RVLM (16±11 spikes/s) neurons.

### Responses to Changes in Head Position

Analyses of data focused on units for which multiple 40° tilts (≥5) were completed, so we could statistically ascertain whether firing rate differed in the nose-down and nose-up positions [9]. A total of 45 LTF units (46% of sampled neurons; 32 with CRA) and 25 NTS units (44% of sampled neurons; 21 with CRA) were included in this statistical analysis, as indicated in Table 1. For the 32 LTF units with CRA, the following number of tilts were performed: 5 for 7 units, 6 for 7 units, 7 for 8 units, 8 for 6 units, 9 for 2 units, 10 for 1 unit, and 11 for 1 unit. For the 21 NTS units with CRA, the following number of tilts were performed: 5 trials for 8 units, 6 trials for 3 units, 7 trials for 3 units, 8 trials for 1 unit, 9 trials for 1 unit, 10 trials for 3 units, 11 trials for 1 unit, and 13 trials for 1 unit. The locations of the LTF and NTS units with CRA are indicated in Fig. 3.

#### Responses to 40°Static Tilts

The activity of units when animals were positioned 20° nose-down was compared using a two-tailed paired t-test to that when animals were positioned 20° nose-up. Fig. 1 illustrates the experimental paradigm, whereas Fig. 5 illustrates the change in activity of an NTS neuron during 10 tilt trials; the activity of the unit when the animal was positioned nose-down was significantly lower than when the animal was nose-up (P<0.0001). Of the 32 LTF units with CRA, the activity of 12 (38%) was significantly different (P<0.05) between the nose-down and nose-up positions. Similarly, the activity of 8 of 21 NTS units with CRA (38%) was significantly different when animals were positioned nose-down and nose-up. Fig. 6 shows the average percent change in firing rate during nose-up rotations for the neurons with significant responses to the change in body position. The firing of the majority of LTF (7/12) and NTS (5/8) neurons was higher when the animals were in the nose-up position. The mean absolute change in firing rate during 40° nose-up tilt was 51±25% for LTF neurons and 51±23% for NTS neurons. These changes in activity for NTS and LTF neurons were not significantly different (P=0.94, unpaired two-tailed t-test). Fig. 7 shows the activity during every trial for the LTF and NTS neurons that responded to tilts; P values over each panel show the significance of the responses. In most cases, the changes in unit activity were similar between trials.

**Fig. 5.**
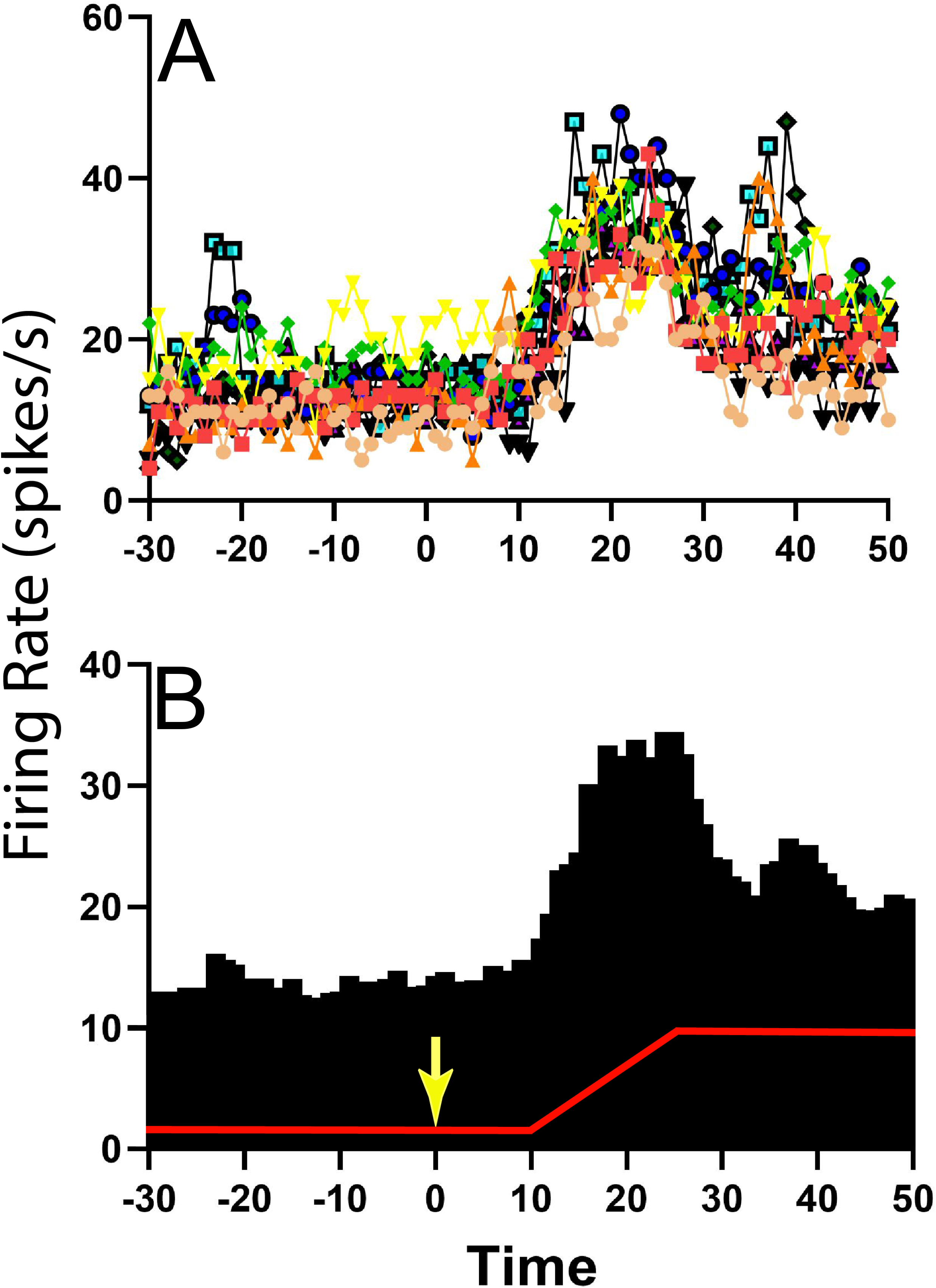
Change in activity for one NTS unit with CRA during 40° head-up tilt. *A:* Unit activity during one second bins for 10 individual trials; each trial is demarked by lines of different colors as well as different symbols. *B:* Histogram of averaged activity during one second bins for all trials. The red line designates table position; the transition from 20° nose-down to the 20° nose-up position (40° change in animal position) occurred 10 seconds after the light cue, which is demarked by a yellow arrow.

**Fig. 6.**
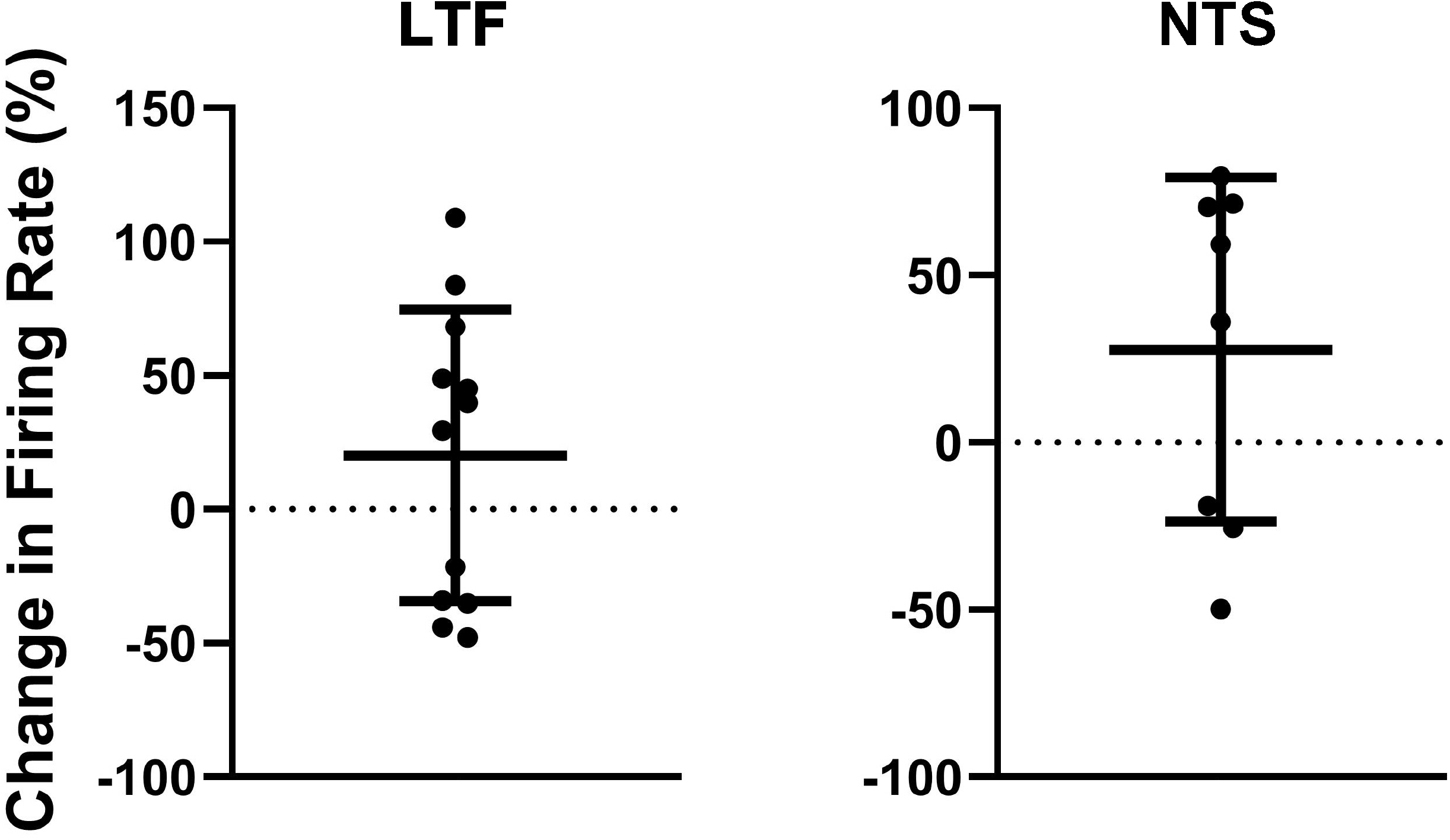
Percent change in firing rate when the animal was tilted nose-up (firing rate in nose-up position ÷ firing rate in the nose-down position) for the 12 units in the LTF and 8 units in the NTS with CRA whose activity was significantly modulated during 40° head-up tilts. Error bars indicate mean ± one standard deviation.

**Fig. 7.**
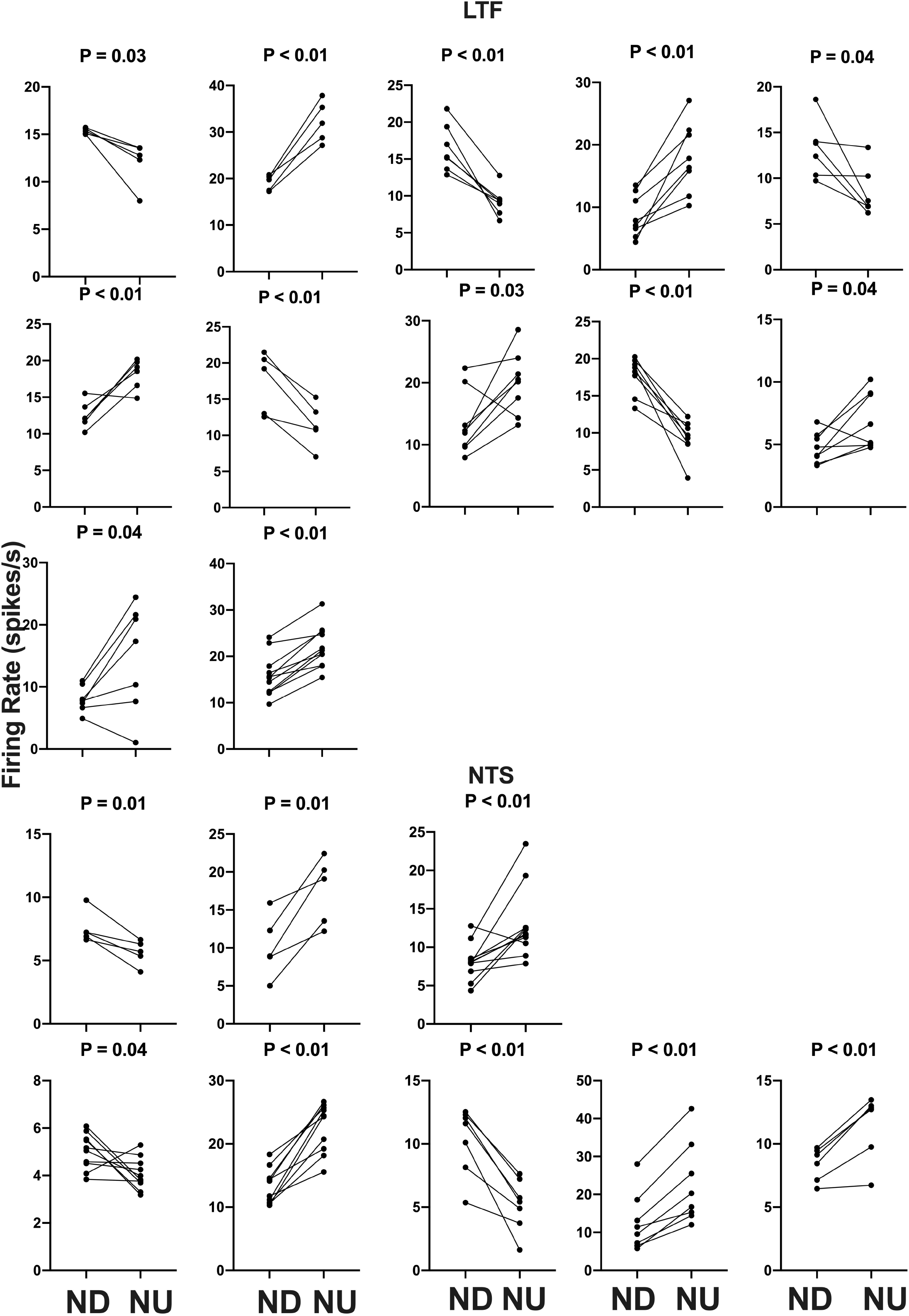
Firing rate during each trial for the 12 LTF and 8 NTS units with CRA whose activity was significantly modulated by 40° head-up tilt. Each panel depicts the firing rate for a single unit when the animal was positioned 20° nose-down (ND) and then subsequently positioned 20° nose-up (NU). P-values indicating the significance of the difference in firing rate in the ND and NU positions (paired t-test) are provided above each panel. The greatest significance level indicated is P<0.01.

LTF and NTS units with CRA whose firing rates were significantly altered by 40° head-up tilt were intermixed with neurons whose firing rates were unaffected by the tilts, as indicated in Fig. 3. LTF neurons that responded to tilts were located on average 2.2±1.3 mm rostral to the obex and 3.0±1.0 mm lateral to the midline, while unresponsive cells were located 2.5±1.2 mm rostral to the obex and 2.6±0.9 mm lateral to the midline. A two-way ANOVA analysis showed that distances from the obex (P=0.74) and midline (P=0.60) were not significantly different for LTF neurons whose activity was significantly modulated or unaffected by the 40° nose-up tilts. NTS neurons that responded to tilts were located on average 1.9±0.56 mm rostral to the obex and 2.7±0.48 mm lateral to the midline, while unresponsive units were located 2.1±0.47 mm rostral to the obex and 2.5±0.5 mm lateral to the midline. A two-way ANOVA analysis revealed that distances from the obex (P=0.37) and midline (P=0.26) were not significantly different for NTS units that responded to or were unaffected by the 40° nose-up tilts.

We also considered whether the firing rates of LTF units lacking CRA were significantly modulated during 40° changes in body position. Out of the 13 LTF cells lacking CRA with five or more tilt trials, seven (54%) responded significantly to 40° head-up tilts. P values ranged from 0.0004 to 0.02, with five responses having P<0.01. Activity decreased in four units and increased in the other three during head-up tilts. The average absolute change in firing rate for the seven units that responded significantly to 40° head-up tilts was 26±21%. A two-sided Fisher’s exact test failed to reveal any significant difference in the proportion of LTF units with and without CRA that responded to tilts (P=0.31). A similar analysis was not conducted for NTS neurons, as the firing rate of only four neurons lacking CRA was recorded during five or more tilts. One of these neurons responded significantly to head-up rotations.

To determine if inputs from baroreceptors might be responsible for responses of LTF and NTS neurons with CRA to 40° head-up tilts, we also considered changes in heart rate during 40° rotations. Heart rate changed significantly (P<0.05) during rotations for only 2/12 (17%) LTF units that responded to 40° tilts and for 5/20 (25%) of the units whose activity was not significantly modulated by 40° tilts. In 6/7 (85%) of these cases, heart rate decreased during head-up tilts, and the magnitude of the heart rate changes was small: 3.5±0.10% for units that responded to tilts and 3.4±5.9% for those without significant responses to the rotations. Significant changes in heart rate occurred during rotations for 3/8 (38%) of the neurons in the NTS that responded to 40° head-up tilts and 3/13 (23%) that failed to respond these movements. Heart rate decreased in all of these cases during head-up tilt. On average, heart rate decreased by 5.4±0.41% for trials in which the firing rates of NTS neurons were modulated by 40° head-up tilts and decreased by 6.0±1.2% for trials in which no significant changes in firing rate occurred during the rotations. The changes in heart rate during tilts that altered or failed to alter the firing rate of cells in the NTS were not significantly different (P=0.32), as indicated by a two-tailed t-test.

#### Responses to 10° Sinusoidal Tilts

Responses were also recorded during 10° sinusoidal rotations in the pitch plane at 0.5 Hz for the 12 LTF units and 8 NTS units that responded to static head-up tilts. Only two of the LTF units (17%) and one unit in the NTS (13%) had significant responses to these rotations, in accordance with established criteria [8; 22; 24; 30; 31]. The responses of two LTF units to 10° sinusoidal rotations are illustrated in Fig. 8. All 3 units that responded significantly to 10° sinusoidal rotations also had large (>30%) changes in activity during 40° head-up tilts.

**Fig. 8.**
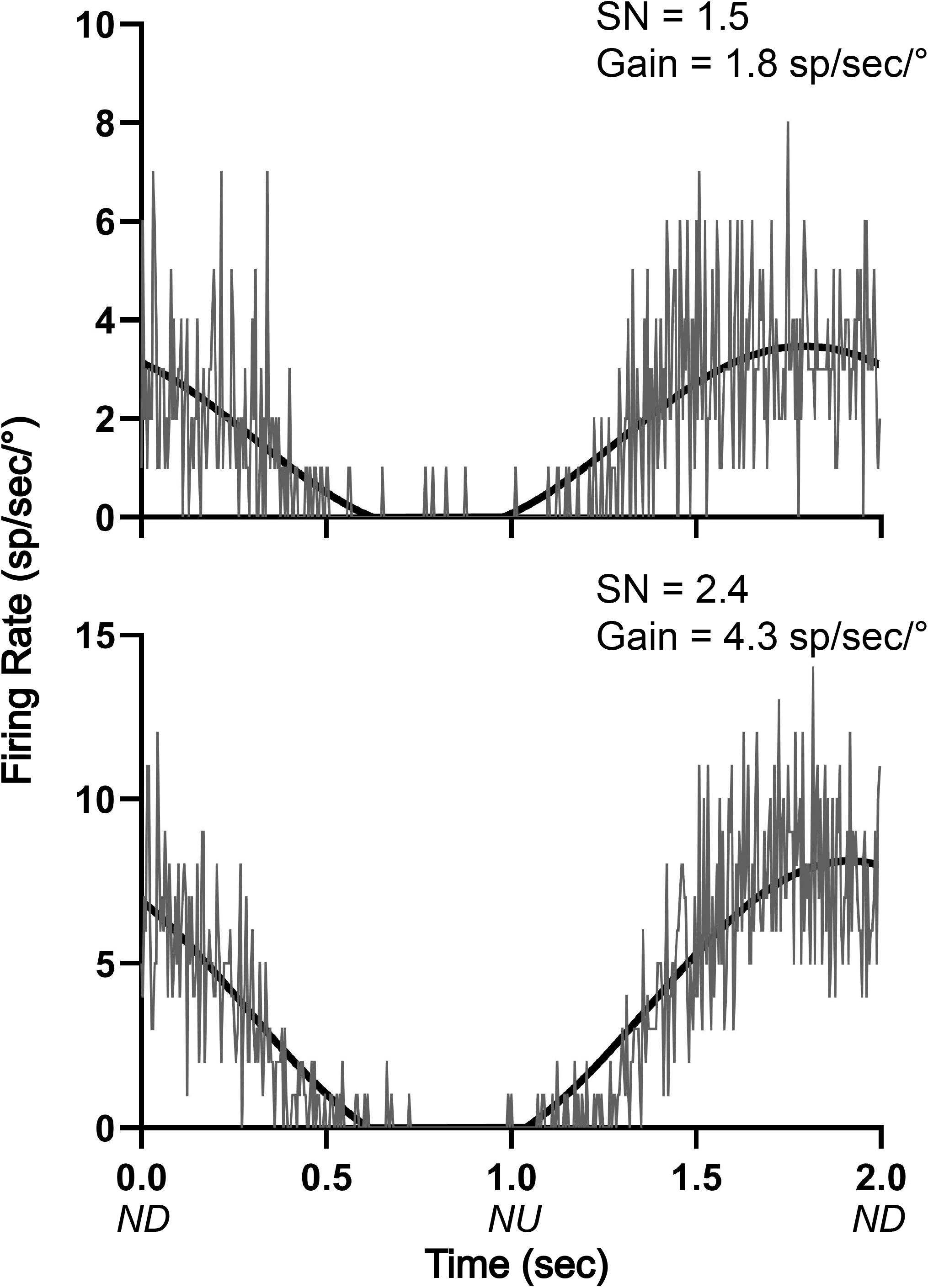
Averaged responses (^~^30 trials) of two LTF units with CRA to 10° sinusoidal rotations in the pitch plane at 0.5 Hz. Bin width was 4 msec (500 bins/trace). A sine wave fitted to responses is shown by black lines. The signal-to-noise (SN) ratio and gain for each unit’s response is indicated. Abbreviations: ND, nose-down; NU, nose-up.

### Responses to Light Cue Preceding Static Tilts

Throughout the course of experimentation, a light cue was provided 10 seconds prior to the onset of each 40° head-up tilt, and we determined whether the activity of LTF or NTS units changed after this cue and before the onset of tilts (see Fig. 1). As indicated in Fig. 9, two-tailed paired t-tests revealed that only 3/32 LTF (9%) units with CRA, and just one that responded to head-up tilt, exhibited significant (P<0.05) changes in activity in the 5-second time period prior to tilt onset (Post-2 period indicated in Fig. 1) compared to the 5-second time period prior the light cue (Pre-period indicated in Fig. 1). Only 1/21 NTS units with CRA, and none that responded to head-up tilts, had a significant increase in firing rate during the Post-2 period. A linear regression analysis failed to show a strong correlation between the significance of responses to the light cue and head-up tilts for LTF (P=0.11, R^2^=0.23) or NTS (P=0.03, R^2^=0.21) neurons.

**Fig. 9.**
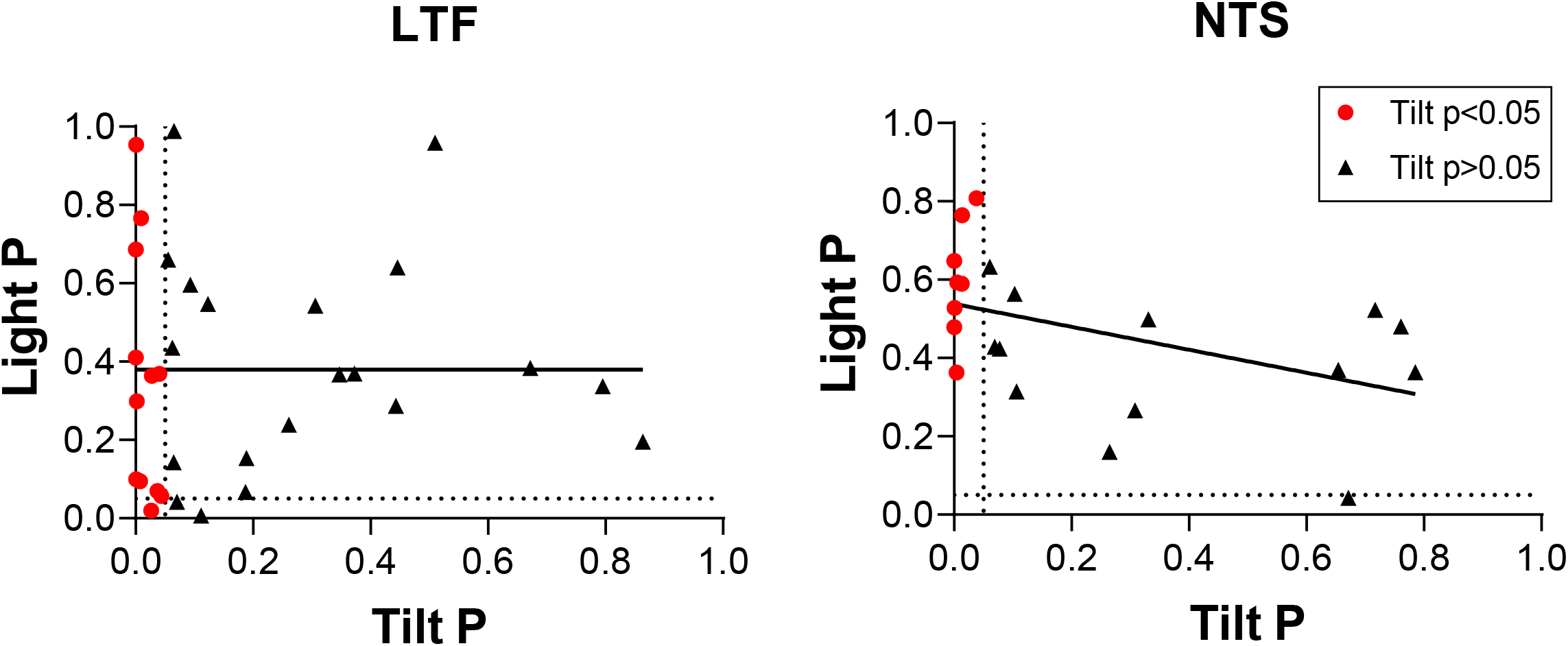
Comparison of P-values (two-tailed t-test) for responses to head-up tilt (Tilt P) and responses to the light cue preceding tilts (Light P, pre-tilt vs post-2 period indicated in Fig. 1). Data for neurons with significant responses (P<0.05) to head-up tilts are indicated by red symbols. Lines show the best-fit of the data with a linear regression analysis.

## 4 Discussion

These experiments included the first recordings in conscious animals from NTS and LTF neurons with changes in firing rate correlated with the cardiac cycle, which presumably participate in regulating cardiovascular responses mediated by the sympathetic nervous system [4]. They showed that the activity of an appreciable fraction (38%) of both NTS and LTF neurons was significantly modulated by 40° static head-up tilts. The firing rates of NTS and LTF neurons with CRA changed on average ^~^50% during the 40° head-up tilts. Such responses are similar to those of RVLM neurons during 40° static head-up tilts: 31% responded to these rotations, with an average change in firing rate of 34% [9]. There was not a significant difference in the magnitude of responses to head-up tilts for NTS, LTF, and RVLM neurons (p=0.1, one-way ANOVA). Since RVLM neurons receive direct and/or multisynaptic inputs from NTS [1; 2; 3] and LTF [14; 15; 16; 17], and the vestibular nuclei project to both regions [18; 19; 20; 21], these data are consistent with the hypothesis that LTF and NTS neurons are components of the vestibulo-sympathetic reflex pathway.

Although the firing rate of over a third of NTS and LTF neurons with CRA was modulated by static head-up tilts, few of these neurons responded to 10° sinusoidal rotations of the body in the pitch plane. Studies in decerebrate animals have shown that an appreciable fraction of NTS neurons with baroreceptor inputs (54%) [22] and LTF neurons (^~^60%) [23] responded to sinusoidal body rotations <10° in magnitude. Thus, as for RVLM neurons [8], LTF and NTS neurons with CRA only responded to large-amplitude body rotations that could produce fluid shifts requiring compensatory changes in sympathetic nervous system activity. In contrast, neurons in caudal regions of the vestibular nuclei that provide inputs to LTF and NTS responded robustly to small-amplitude body rotations in conscious felines [30; 32]. Considering that the caudal aspect of the vestibular nuclei has direct projections to NTS and LTF [18; 19; 20; 21], it is unclear how gating of labyrinthine signals occurs in vestibulo-sympathetic responses, such that sympathetic nerve activity only changes during large-amplitude body rotations. It is feasible that NTS and LTF neurons also receive convergent inputs from other regions of the nervous system that modify their responses to vestibular stimuli. It has been postulated that the caudal cerebellar uvula modulates vestibulo-sympathetic responses [10], and the present data raise the prospect that a pathway originating in the cerebellum or elsewhere in the nervous system alters the responsiveness of LTF and NTS neurons to vestibular stimuli. This hypothesis remains to be validated experimentally.

We also evaluated whether the responses of NTS and LTF neurons to large-amplitude static tilts could be due to activation of the baroreceptor reflex. Heart rate changes related to tilts were small and significant during trials for only a small proportion of neurons with CRA whose neuronal activity changed appreciably during the rotations. In addition, significant heart rate changes were noted for approximately equal proportions of cells whose firing rates were and were not modulated by head-up rotations. These data suggest that the primary signals altering the firing rates of NTS and LTF neurons during whole-body tilts were not from baroreceptors.

The magnitude of CRA of LTF and NTS neurons, gauged by the ratio of peak to background counts time-locked to the cardiac cycle, varied between trials. However, this variability was significantly smaller than for RVLM neurons [9]. This may be a reflection of the complexity of inputs to the RVLM, which include projections from a variety of brainstem and supratentorial structures [4]. These data suggest that factors such as alertness and vigilance have less of an impact on the excitability of LTF and NTS neurons than RVLM neurons, although this prospect is yet to be examined experimentally.

A light cue preceded 40° head-up tilts in these experiments, but there was no evidence that the activity of LTF or NTS units changed after the light cue and prior to the whole-body rotations. A previous experiment provided similar results for RVLM neurons [9]. An earlier study showed that heart rate and cerebral blood flow was unaffected by a light cue delivered before 60° head-up tilts in conscious felines [28]. Collectively, these data provide strong evidence that feedforward (preparatory) cardiovascular responses are absent prior to imposed postural changes that can result in a change in blood distribution in the body. This is in contrast to exercise and some other active movements (or imagination of active movements) in which cardiovascular responses occur prior to movement onset [25; 26; 27]. There is considerable evidence that motor cortex has multisynaptic influences on sympathetic nervous system activity [33; 34; 35], as do subcortical areas involved in motor control [27]. One possibility is that feedforward cardiovascular responses prior to movement are elicited through connections of brain motor areas with RVLM neurons, such that anticipation of imposed movements that do not entail motor responses fail to trigger changes in heart rate or peripheral blood flow. Additional experiments are needed to examine this hypothesis.

LTF and NTS neurons with and without CRA had similar responses during whole-body rotations in these studies. Studies in decerebrate and anesthetized animals have shown that neurons in these regions with respiratory-related activity and inputs from gastrointestinal receptors, as well as those with baroreceptor inputs, responded to vestibular stimuli [19; 22; 23]. Head-up body rotations require changes in the activity of respiratory muscles to compensate for gravitational loading on these muscles [36], and both LTF and NTS neurons have been proposed as components of brainstem circuity that produces motion sickness [37]. It is thus not surprising that NTS and LTF neurons lacking CRA responded to vestibular stimuli. However, the design of these experiments did not permit a determination of the physiologic role of neurons lacking CRA.

In summary, these experiments included the first recordings in conscious animals from NTS and LTF neurons with CRA that presumably participate in regulating cardiovascular responses mediated through the sympathetic nervous system. The studies showed that the responses of these neurons to whole-body rotations that activate the vestibular system were similar to those documented for RVLM neurons [9]. In combination with other findings, these data support the conclusion that LTF and NTS neurons are components of the brainstem pathway that mediates vestibulo-sympathetic responses. However, variability in CRA from trial to trial was smaller for NTS and LTF neurons than for RVLM neurons. This observation suggests that RVLM neurons may receive stronger inputs from supratentorial regions than those in NTS and LTF, such that their excitability is modified more powerfully in accordance with an animal’s vigilance and alertness.

## Acknowledgments

The authors thank Rutha Chivate, Asmita Joshi, Emmanuel Kambouroglos, Robert Khorami, John Lausch, Christian Molfetto, Bridget Perry, Sophie Tayade, Anisha Venkatesh, and Samuel Wittman for their assistance with data collection and analysis.

